# Hierarchical Bayesian Modelling Improves Microstructural Parameter Mapping in Diffusion and Exchange MRI Data

**DOI:** 10.1101/2025.09.04.674046

**Authors:** Elizabeth Powell, Mark Maskery, Hedley C.A. Emsley, Laura M. Parkes, Geoff J.M. Parker, Paddy J. Slator

**Affiliations:** Department of Medical Physics and Biomedical Engineering, University College London, London, UK; Lancaster Medical School, Lancaster University, Lancaster, UK; Department of Neurology, Lancashire Teaching Hospitals NHS Foundation Trust, Preston, UK; Division of Psychology, Communication and Human Neuroscience, School of Health Sciences, Faculty of Biology, Medicine and Health, University of Manchester, Manchester, UK; Geoffrey Jefferson Brain Research Centre, Faculty of Biology, Medicine and Health, University of Manchester, Manchester, UK; Bioxydyn Limited, Manchester, UK; Cardiff University Brain Research Imaging Centre, School of Psychology, Cardiff University, Cardiff, UK; School of Computer Science and Informatics, Cardiff University, Cardiff, UK

**Keywords:** Bayesian modelling, blood-brain barrier, diffusion MRI, filter exchange imaging, kurtosis, microstructure, water exchange

## Abstract

Microstructure modelling quantifies subvoxel tissue features by combining an MRI acquisition with a mathematical model, which is typically fitted voxel-by-voxel with least-squares (LSQ) minimisation to give voxelwise maps of microstructural quantities such as diffusivity and compartmental fractions. Such approaches are susceptible to voxelwise noise, which can lead to erroneous values in parameter maps. Hierarchical Bayesian modelling (HBM) can address this limitation, but has only been demonstrated for simple models.

We previously derived an HBM approach for an arbitrary microstructure model with flexible parameter constraints, utilising a Markov chain Monte Carlo algorithm for parameter estimation; here the method is demonstrated and evaluated using simulated and human data for two previously unexplored diffusion MRI techniques, namely diffusion kurtosis imaging and blood-brain barrier filter exchange imaging. When compared with LSQ minimisation, HBM increased the accuracy, precision, contrast-to-noise ratio, and parameter map quality in both simulated and human data. HBM was also able to resolve local parameter variations associated with white matter lesions in a small sample of cerebral small vessel disease subjects, which were obscured by high noise levels in the LSQ-derived parameter maps. Finally, a noise sensitivity assessment in simulations showed that HBM improved the contrast-to-noise ratio and parameter map quality even at low signal-to-noise ratios.

This generalised HBM framework can improve parameter estimation for more complex diffusion MRI microstructural models that extend beyond linear combinations of exponentials.

## 1 INTRODUCTION

Diffusion MRI (dMRI) is sensitive to the microscopic motion of water molecules. Microstructure imaging combines specialised dMRI acquisitions with tissue models to estimate quantitative parameters of tissue microstructure and microcirculation. This approach has been widely applied in neuroimaging - demonstrating estimation of neurite density ^1^, soma density ^2^, axonal diameter ^3^, and water exchange ^4–6^ - as well as being used in body imaging ^7–9^.

Such tissue models are typically fitted to dMRI data in each voxel separately to estimate the model parameters. This process has traditionally employed least squares (LSQ) regression, although machine learning is emerging as an attractive alternative that can improve the precision ^10^ of microstructural maps. However, maps generated through voxel-by-voxel fitting are heavily affected by noise in individual voxels and do not optimally exploit the data, disregarding dependencies between voxels. In supervised machine learning approaches, the distribution of the training data effectively acts as a prior on the parameter estimates, providing some degree of noise-robustness; however, this prior is fixed in advance by the choice of training data, and it has been shown that this choice influences the resulting estimates ^11^. Methods that explicitly capture dependencies between neighbouring voxels demonstrate fast and robust fits - often using convolutional neural networks ^12–15^ - but these spatial regularisation techniques only capture patterns across predefined local windows, rather than capturing regional or global patterns.

Hierarchical Bayesian modelling (HBM) offers a promising alternative approach that breaks the assumption of independent voxels, whilst allowing for the incorporation of dependencies at various scales to enhance fitted maps ^16^. HBM assumes a prior, typically Gaussian in this context, on model parameters over a region of interest (ROI). Rather than being user defined, the prior is estimated in a data-driven manner. This prior reduces sensitivity to noise in single voxels, leading to improved parameter maps. Orton et al. ^16^ introduced the HBM approach for diffusion MRI, formulating the model and a Markov chain Monte Carlo (MCMC) algorithm for fitting the intravoxel incohorent motion (IVIM) model to dMRI data, and demonstrated improved IVIM parameter maps in the liver. However, HBM has still primarily only been applied to a limited number of simple models. Notably, it has been used extensively with the IVIM model ^16–26^, as well as with a three compartment model for placental MRI ^27^.

In this paper, we build on our previous preliminary work ^28^ and demonstrate our general (i.e. non model specific) hierarchical Bayesian framework for microstructure modelling in two advanced neuroimaging applications. The extension constitutes: (i) evaluation of the framework in both simulated and in vivo data in two previously unexplored neuroimaging applications, namely mean signal diffusion kurtosis imaging (DKI) ^29^, and blood-brain barrier filter exchange imaging (BBB-FEXI) ^5,6^ using the apparent exchange rate (AXR) model ^4^; (ii) an assessment of the noise sensitivity of the regional priors; (iii) an analysis of the ability of different regional priors to resolve local parameter variations (e.g. lesions); (iv) a full derivation of the method, given in Appendix A for completeness, and; (v) presentation of two open source toolboxes, one Matlab (https://github.com/e-powell/matlab-bayesian) and one Python (https://github.com/e-powell/dmipy-bayesian). The HBM approach is shown to enhance parameter mapping relative to traditional voxel-by-voxel fitting, demonstrating that the advantages of HBM in dMRI extend beyond models that are linear combinations of exponentials.

## 2 METHODS

### 2.1 Background: a general hierarchical Bayesian microstructure model

The mathematical formulation for a general hierarchical Bayesian microstructural modelling framework is summarised here, building on our preliminary work ^28^ which extends the methods of Orton et al. ^16^; see Appendix A for the full derivation and algorithm pseudocode.

Consider a general microstructure model (Appendix A.1) with *C* compartments and a set of microstructure-related parameters given by:

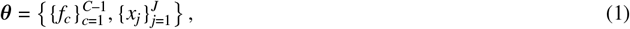

where *f*_*c*_ denotes compartment signal fractions and *x*_*j*_ all other parameters (e.g. diffusivities, orientations, radii). The signal equation in voxel *i* for an acquisition of length *N* can be written:

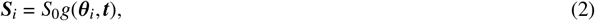

where ***S***_*i*_ = [*S*_1_, …, *S*_*N*_]^*T*^ is the noise-free model-predicted signal, ***t*** = [*t*_1_, …, *t*_*N*_]^*T*^ is the set of acquisition parameters (e.g. diffusion encodings), and ***θ***_*i*_ represents the underlying parameter values.

A Bayesian shrinkage prior (BSP) is assumed, following Orton et al. ^16^, but is here extended to accommodate multiple ROIs (Appendix A.2). The prior on parameters in voxel *i*, which is in ROI *k*, is therefore:

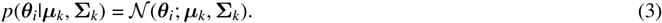

The posterior distribution and BSP parameters over each ROI, ***µ***_*k*_ and **Σ**_*k*_, can be directly formulated ^16^. The posterior distribution (Appendix A.3) is:

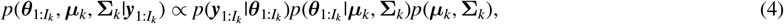

where 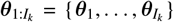 are the microstructure parameters and 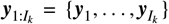 the measured dMRI signal for all *I*_*k*_ voxels in ROI *k*, where each ***y***_*i*_ = [*y*_1_, …, *y*_*N*_]^*T*^ corresponds to ***S***_*i*_ with added noise. Samples can be drawn from this posterior distribution with an MCMC algorithm. Once more following Orton et al. ^16^, the updates for ***µ***_*k*_, **Σ**_*k*_ are Gibbs moves (Appendix A.5.1) and the updates for 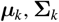 are Metropolis-Hastings moves (Appendix A.5.2). Single parameter estimates for each voxel are obtained by taking the mean of the posterior distribution after the MCMC burn-in period.

A schematic of the full HBM framework is shown in Figure 1. The framework is demonstrated here in two distinct microstructure estimation problems: (i) the well-known mean signal DKI ^29^ experiment, and; (ii) the more recent innovation of filter exchange imaging (FEXI) ^4^ applied to the blood-brain barrier (BBB) ^5,6^.

**FIGURE 1.**
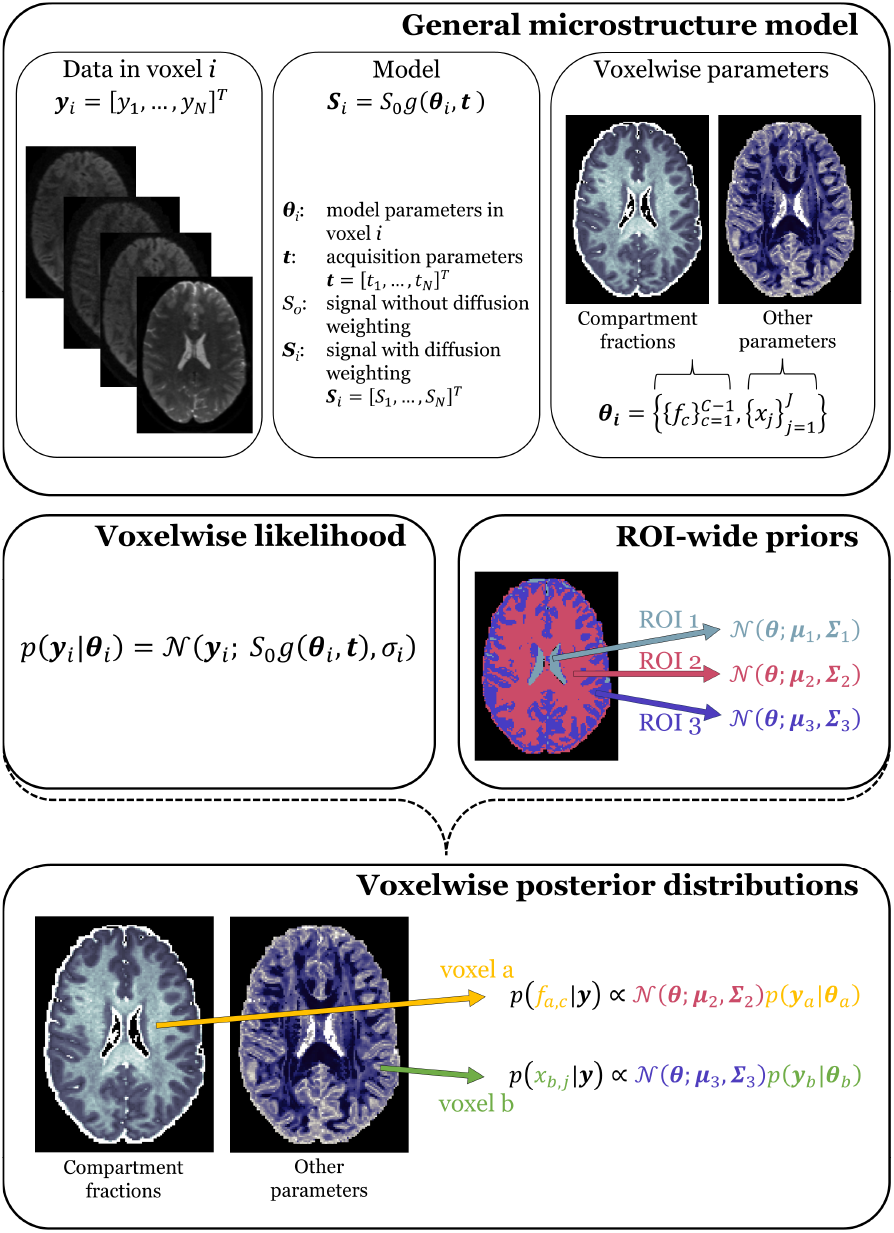
Schematic of the hierarchical Bayesian modelling method. **Top panel**. A general microstructure model, *g* (***θ***_*i*_, ***t***), maps the microstructure-related model parameters ***θ***_*i*_ and the diffusion MRI acquisition parameters ***t*** to the signal ***S***_*i*_ in voxel *i*, where the acquisition volume in a multi-volume acquisition is indexed *n* = 1, …, *N*. The model parameters can be grouped according to parameter type as 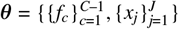, where 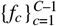 are the compartmental signal fractions and 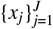 represent all other parameters. **Middle panel**. Definitions of the voxelwise likelihood function and ROI-wide Gaussian priors. **Bottom panel**. Definitions of the voxelwise parameter posteriors, and examples of the corresponding parameter maps.

### 2.2 Exemplar microstructure models

#### 2.2.1 Diffusion kurtosis imaging

DKI ^29^, an extension of diffusion tensor imaging (DTI), provides information on the non-Gaussianity of water diffusion in the brain, from which microstructural tissue information can be inferred. DKI modelling uses higher order cumulant expansion terms of the displacement distribution to extract kurtosis information. This requires higher *b*-values than needed for DTI, which, in addition to the increased model complexity, subsequently increases noise susceptibility and commonly leads to extreme values in the parameter maps when using conventional estimation methods.

The spherically-averaged DKI signal model is given by ^30^:

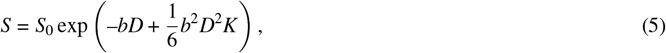

where *S*_0_ is the signal without diffusion encoding, *D* is the apparent diffusion coefficient, and *K* the kurtosis.

#### 2.2.2 Blood-brain barrier filter exchange imaging

Models parameterising water exchange rates across the blood-brain barrier (BBB) are highly noise-sensitive, often requiring spatial or cross-subject data averaging ^5,6,31–33^. Bayesian techniques offer a potential solution to this problem, and could enable subject-specific voxel-level exchange rate estimates.

The BBB-FEXI technique ^5,6^ utilises a double diffusion encoding sequence that was originally developed to measure the apparent exchange rate (AXR) of water across cell membranes ^4,34^. First, a low *b*-value diffusion filter selectively suppresses the signal from fast pseudo-diffusing intra-vascular spins. The intra-vascular signal recovers during the mixing time via exchange across the BBB, such that the signal measured after the second diffusion encoding block is a weighted combination of intra- and extra-vascular signals, dependent on the mixing time and exchange rate. The FEXI AXR model ^4^ is defined as:

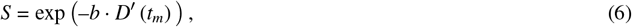

where

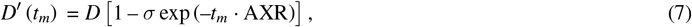

and *D* and *D*^*′*^ (*t*_*m*_) are the apparent diffusion coefficients at equilibrium (i.e. filter inactive) and at each mixing time *t*_*m*_ with the filter active, respectively, and *σ* is the filter efficiency.

### 2.3 Data

The HBM algorithm is demonstrated in both the DKI and FEXI AXR models first in simulations, where there is a known ground truth, before being applied in human data.

#### 2.3.1 Diffusion kurtosis imaging

##### Simulations

Synthetic data were generated for two distinct ROIs representative of healthy human white matter (WM) and grey matter (GM). Parameter values were obtained by fitting the DKI model (Equation 5) using LSQ estimation to a randomly selected subject from the Human Connectome Project (HCP; details below) and calculating the mean and standard deviation (SD) of fitted parameters in WM/GM. Ground truth parameters were then drawn from normal distributions defined over these means and SDs: in WM, they were *D*_*wm*_ *∼ N*(0.87, 0.29) µm^2^/ms, *K*_*wm*_ *∼ N*(1.04, 0.41); in GM they were *D*_*gm*_ *∼ N*(1.12, 0.50) µm^2^/ms, *K*_*gm*_ *∼ N*(0.63, 0.22). The total matrix size was 50 *×* 50, with 1204 voxels in the ‘WM’ ROI and 1296 in the ‘GM’ ROI. Zero-mean Gaussian noise was added to the complex signal (i.e. to the real and imaginary components independently) to give a signal-to-noise ratio SNR = 20 in the *b* = 0 data. Signals were generated for *b* = 0, 1, 2, 3 ms/µm^2^ with a number of signal averages NSA = 2, 9, 9, 9 respectively, mimicking multiple gradient directions (Table 1).

**TABLE 1.** DKI protocol (HCP). The DKI parameters shown are those used by the Human Connectom Project (HCP). Note that only 2,9,9,9 gradient directions (29 volumes total) were used for the analysis here. The T_1_-weighted MPRAGE data (resolution = 0.7 × 0.7 × 0.7 mm^3^; TR = 2400 ms; TE = 2.14 ms; flip angle = 8 ^°^) were additionally utilised to obtain WM/GM segmentations. **BBB-FEXI protocol (cSVD)**. Cerebral small vessel disease (cSVD) subject data were acquired on a Philips 3 T Ingenia Elition X system using an in-house double-diffusion encoding sequence. A T1-weighted FFE (resolution = 1×1×1 mm^3^; TR = 6.8 ms; TE = 3.1 ms; flip angle = 8 ^°^) was acquired for WM/GM segmentation, and a FLAIR (resolution = 0.86 × 0.86 × 1 mm^3^; TR = 4800 ms; TE = 340 ms; inversion time TI = 1650 ms) for WM hyper-intensity segmentation.

|  | DKI (HCP) | BBB-FEXI (cSVD) |  |  |  |
| --- | --- | --- | --- | --- | --- |
| Resolution [$\text{mm}^3$] | $1.25 \times 1.25 \times 1.25$ | $3 \times 3 \times 5$ | | | |
| No. slices | 111 | 18 |  |  |  |
| Repetition time, TR [ms] | 5520 | 5000 |  |  |  |
| Echo time, TE [ms] | 90 | 63 |  |  |  |
| Filter block echo time, $TE_f$ [ms] | - | 42 | | | |
| $b$-values, $b$ [$\text{ms}/\mu\text{m}^2$] | 0, 1, 2, 3 | 0, 0.25 | 0, 0.25 | 0, 0.25 | 0, 0.25 |
| Gradient directions | 18, 90, 90, 90 | 6, 6 | 6, 6 | 6, 6 | 6, 6 |
| No. repetitions, $N_{reps}$ | 1 | 1, 1 | 1, 1 | 1, 1 | 1, 1 |
| Mixing time, $t_m$ [ms] | - | 16 | 16 | 200 | 400 |
| Filter block $b$-values, $b_f$ [$\text{s}/\text{mm}^2$] | - | 0 | 0.25 | 0.25 | 0.25 |
| Total no. volumes | 288 | 48 |  |  |  |

**Abbreviations:** BBB-FEXI, blood-brain barrier filter exchange imaging; DKI, diffusion kurtosis imaging; FFE, fast field echo; FLAIR, fluid-attenuated inversion recovery; MPRAGE, magnetization-prepared rapid acquisition gradient echo.

##### Human Connectom Project data

Images for two subjects were obtained from the publicly-available data sets provided by the HCP ^35^ (WU-Minn Consortium; 1U54MH091657; funded by the 16 NIH Institutes and Centers that support the NIH Blueprint for Neuroscience Research and by the McDonnell Center for Systems Neuroscience at Washington University) (Test Retest Data Release, release date: Mar 01, 2017, available online at humanconnectome.org). To better demonstrate the efficacy of the HBM approach, which is most beneficial when applied to lower SNR data, the number of gradient directions was reduced to 2, 9, 9, 9 for *b* = 0, 1, 2, 3 ms/µm^2^ respectively, in line with the NSA of the simulated data. WM/GM segmentations were obtained by first fitting the diffusion tensor then calculating apparent diffusion coefficient (ADC) and fractional anisotropy (FA) maps with the MRtrix3^36^ functions *dwi2tensor* and *tensor2metric* respectively. GM masks were defined by selecting voxels with ADC < 2 µm^2^/ms and FA < 0.15, and WM masks by selecting voxels with ADC < 2 µm^2^/ms and FA > 0.15.

#### 2.3.2 Blood-brain barrier filter exchange imaging

##### Simulations

Model parameters were again generated for two ROIs representing healthy WM/GM with a total matrix size of 50 *×* 50. Parameter values were drawn from normal distributions with means and SDs taken from literature values for healthy subjects ^6^: in the ‘WM’ ROI, these were *D*_*wm*_ *∼ N* (0.90, 0.04) µm^2^/ms, *σ*_*wm*_ *∼ N* (0.14, 0.02), AXR_*wm*_ *∼ N* (2.00, 0.30) s^−1^; in the ‘GM’ ROI they were *D*_*gm*_ *∼ N* (1.20, 0.08) µm^2^/ms, *σ*_*gm*_ *∼ N* (0.18, 0.03), AXR_*gm*_ *∼ N* (1.40, 0.35) s^−1^. Six signals were generated for each parameter combination using the acquisition parameters in Table 1; zero-mean Gaussian noise was added to each complex signal to give SNR = 30 in the equilibrium data (i.e. with the filter inactive, *b*_*f*_ = 0 ms/µm^2^, the minimum mixing time, *t*_*m*_ = 16 ms, and without diffusion weighting, *b* = 0 ms/µm^2^), comparable to the human data (see below and Table 1). Each set of six noisy signals were then averaged (i.e. NSA = 6) to mimic the six gradient directions used in the human data.

##### Cerebral small vessel disease subjects

Two exemplar subjects from an ongoing cerebral small vessel disease (cSVD) study were analysed to provide proof-of-principle in the presence of pathology. Patients were recruited from Lancashire Teaching Hospitals NHS Foundation Trust and Salford Royal NHS Foundation Trust. The study was approved by Yorkshire and The Humber (Leeds West) Research Ethics Committee (21/YH/0119), the Health Research Authority and local research governance panels.

Subjects were scanned on a Philips 3 T Ingenia Elition X system using a 32-channel head coil. BBB-FEXI, T_1_-weighted, and fluid attenuated inversion recovery (FLAIR) data were acquired; see Table 1 for protocols. Susceptibility and eddy current distortions in the BBB-FEXI data were corrected using FSL *topup* and *eddy* ^37,38^. The T_1_-weighted data were segmented using FreeSurfer ^39^ to produce WM/GM ROIs. The T_1_-weighted and FLAIR data were used to produce WM hyper-intensity (WMH) masks using the Lesion Segmentation Toolbox ^40^ in SPM12 with a locally established threshold of 0.3. The T_1_-weighted and FLAIR images were registered to the BBB-FEXI data using FSL FLIRT ^41^, and the respective warps used to propagate the WM/GM/WMH masks to the BBB-FEXI data.

### 2.4 Model fitting

#### 2.4.1 Least-squares estimation

LSQ fitting was performed on all data sets to provide priors for the HBM method and to provide conventional fitting results against which the HBM results could be compared. The LSQ algorithm (*fminsearchbnd* in Matlab R2024b) was initialised using 25 and 27 starting values for the DKI and FEXI AXR models respectively, uniformly distributed between parameter constraints to avoid local minima. The DKI model fits were constrained such that *D* ∈ (0.1, 3.5) µm^2^/ms and *K* ∈ (0, 3); parameter constraints for the FEXI AXR model were *D* ∈ (0.1, 3.5) µm^2^/ms, *σ* ∈ (0, 1), AXR ∈ (0, 50) s^−1^.

#### 2.4.2 Hierarchical Bayesian estimation

The HBM framework implemented in Matlab (R2024b) was used for all data. Parameter values were initialised with the voxelwise LSQ fit and constrained with the same bounds (Section 2.4.1). For the simulated and human DKI data the MCMC algorithm was run for *N*_*s*_ = 1 *×* 10^5^ steps. For the human BBB-FEXI data the MCMC algorithm was similarly run with *N*_*s*_ = 1 *×* 10^5^ steps; for the simulated FEXI AXR data the MCMC chain length was increased to *N*_*s*_ = 4 *×* 10^5^ steps to improve the conditioning of the regional priors given the reduced number of voxels (2500 voxels) compared to the human data (*∼*20,000 voxels). In all cases the burn-in period was defined as the first *N*_*burn*_ = *N*_*s*_/2 steps. Weights were updated every 100 steps during the first half of the burn-in period (Appendix A.5.3), which was heuristically found to be sufficient to sample the posterior distributions. Two ROIs were used for the regional priors - one each for WM/GM - for the DKI data (simulated and human) and the simulated FEXI AXR data; a third ROI was utilised in cSVD patient BBB-FEXI data for WMH.

Parameter posterior distributions and representative statistics were calculated from the MCMC samples after the burn-in. Voxelwise parameter maps were generated using the mean of the posterior distributions in each voxel *i*:

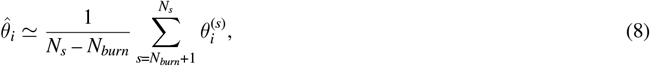

where 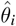 is the expected value of a single parameter (Equation 1) and 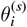 the value sampled at each MCMC step. Summary statistics for the regional priors were calculated as:

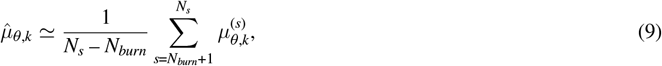

where 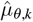 is the prior mean of parameter *θ* in the *k*th ROI and 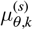 the prior mean estimated at each MCMC step.

### 2.5 Evaluation

A series of metrics were used to test the ability of the methods to: (i) accurately and precisely recover ground truth parameter values under variable SNR conditions (simulations), and; (ii) obtain contrast between different ROIs (simulations and human data).

#### 2.5.1 Accuracy and precision

In simulations, the precision of fitted parameters was evaluated using the root mean squared error (RMSE):

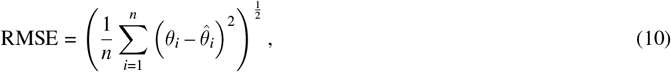

where *θ*_*i*_ is the ground truth of a single model parameter in voxel *i* and 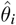 the estimated value. Bias (accuracy) was defined as:

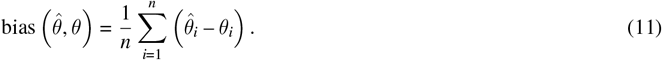

Maps of the percent relative error between ground truth and fitted values were also generated, with the error for voxel *i* given by:

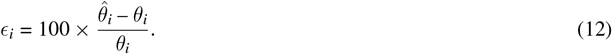

#### 2.5.2 Contrast-to-noise

The contrast-to-noise ratio (CNR) between two ROIs in parameter maps was quantified according to ^42^:

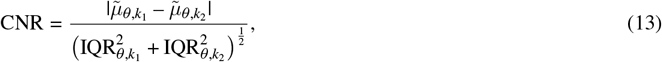

where 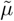 and IQR are the median and inter-quartile range of voxelwise parameter estimates in each ROI.

#### 2.5.3 Noise sensitivity of regional priors

The noise sensitivity of the regional priors was investigated in the DKI and FEXI AXR simulated data. Data were simulated with additional noise characteristics to give three data sets per model: for the DKI model, data were simulated with SNR_*dki*_ = 10, 20, 30; for the FEXI AXR model, data were simulated with SNR_*axr*_ = 10, 30, 50.

#### 2.5.4 Resolving local parameter variations

The ability of the HBM algorithm, and its dependence on the defined regional priors, to resolve small local variations in AXR - as is hypothesised to occur in WMH regions in cSVD subjects - was investigated in the human data. The HBM model was run three times using different prior configurations: (i) one whole brain regional prior (*k* = 1); (ii) two regional priors, one each for WM/GM (*k* = 2), and; (iii) three regional priors, with an additional prior for WMH (*k* = 3). The CNR between WM and WMH was evaluated as a surrogate for lesion discrimination quality for the LSQ output and for each HBM output.

## 3 RESULTS

### 3.1 Accuracy and precision

Figure 2 shows example parameter maps for the simulated data (with SNR_*dki*_ = 20 and SNR_*axr*_ = 30), estimated using the LSQ and HBM approaches. Qualitatively, HBM parameter maps were less affected by noise and showed improved parameter estimation over LSQ maps. The effect of this noise on parameter accuracy and precision is visualised in Figure 3, which shows percent error maps and correlations with ground truth values. Improvements in the estimation of kurtosis and AXR - the most challenging parameters to fit in the DKI and FEXI AXR models respectively - were most apparent: the number of extreme values in the LSQ kurtosis fit was reduced in the HBM fit (Figure 3A; see also Table 2), while the large and numerous errors in the LSQ FEXI AXR outputs were substantially reduced using the HBM approach (Figure 3B). Correlations with ground truth values were also improved using the HBM method: the correlation coefficient increased from *R*_*lsq*_ = 0.87 to *R*_*hbm*_ = 0.93 for kurtosis (Figure 3C) and from *R*_*lsq*_ = 0.14 to *R*_*hbm*_ = 0.41 for AXR (Figure 3D).

**TABLE 2.** Error metrics for the simulated DKI (SNR = 20) and FEXI AXR (SNR = 30) data. Extreme fits are defined as parameter values within 1% of the fitting bounds; values in brackets indicate the total proportion of voxels containing one or more parameters with extreme fits.

|  |  | DKI model |  | AXR model |  |  |
| --- | --- | --- | --- | --- | --- | --- |
| | | $D$<br>[$\mu\text{m}^2/\text{ms}$] | $K$<br>[a.u.] | $D$<br>[$\mu\text{m}^2/\text{ms}$] | $\sigma$<br>[a.u.] | AXR<br>[$\text{s}^{-1}$] |
| **RMSE** | LSQ | $6.5 \times 10^{-2}$ | 0.20 | $8.1 \times 10^{-2}$ | $13.2 \times 10^{-2}$ | 19.2 |
| | HBM | $7.9 \times 10^{-2}$ | 0.15 | $4.6 \times 10^{-2}$ | $3.5 \times 10^{-2}$ | 2.1 |
| **Bias** | LSQ | $0.1 \times 10^{-2}$ | 0.03 | $0.6 \times 10^{-2}$ | $4.3 \times 10^{-2}$ | 8.5 |
| | HBM | $0.2 \times 10^{-2}$ | 0.05 | $0.1 \times 10^{-2}$ | $0.6 \times 10^{-2}$ | 1.0 |
| **CNR** | GT | 0.3 | 0.7 | 2.6 | 0.8 | 1.0 |
|  | LSQ | 0.3 | 0.6 | 1.5 | 0.1 | 0.1 |
|  | HBM | 0.3 | 0.7 | 2.8 | 0.6 | 0.6 |
| **Extreme fits [%]** | LSQ | 0.0 | 2.6 (2.6) | 0.0 | 7.4 | 32.9 (39.6) |
|  | HBM | 0.0 | 0.0 (0.0) | 0.0 | 0.0 | 0.0 (0.0) |

**Abbreviations:** AXR, apparent exchange rate; CNR, contrast-to-noise ratio; DKI, diffusion kurtosis imaging; RMSE, root mean squared error.

**FIGURE 2.**
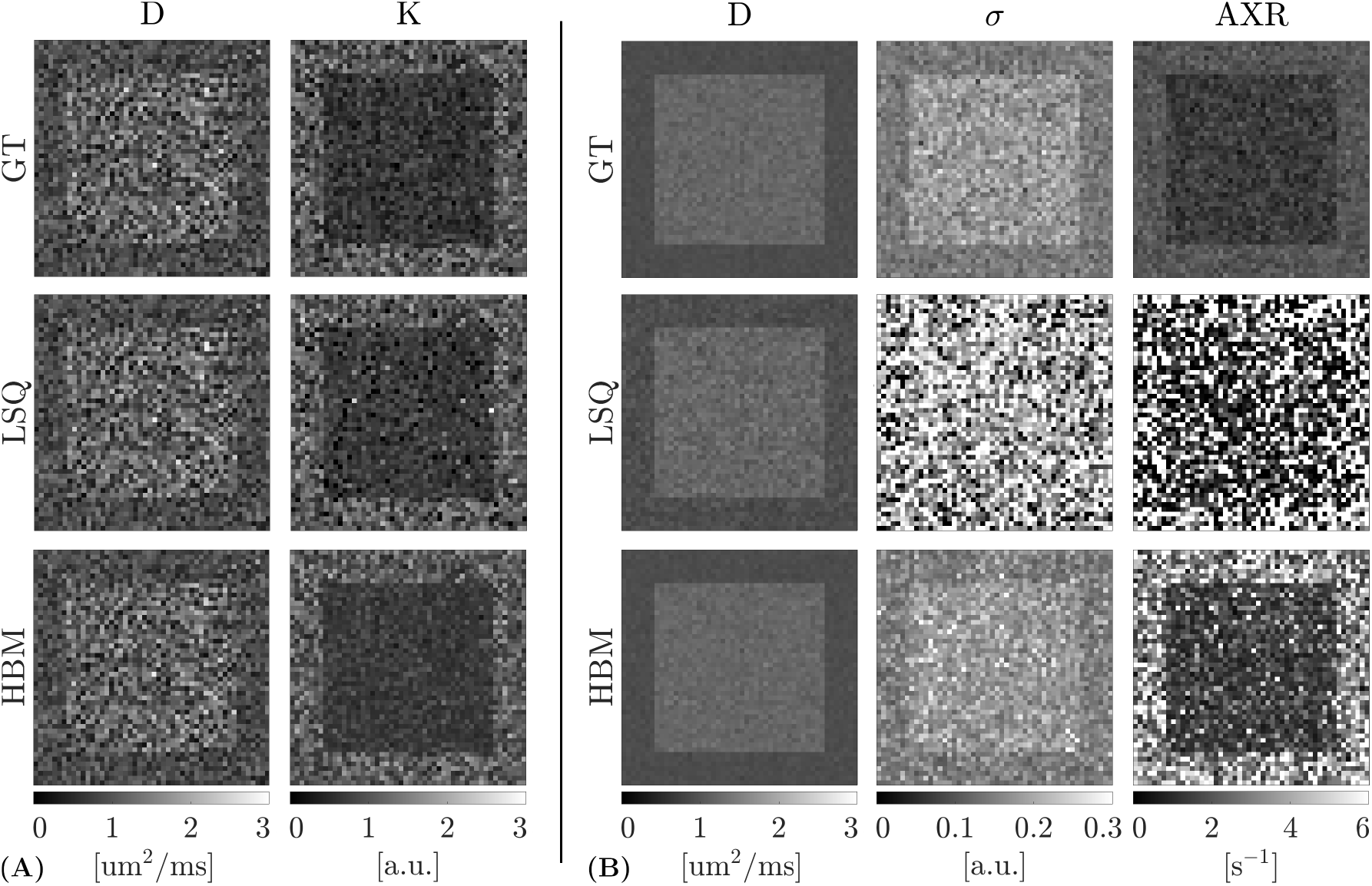
Parameter maps for the simulated data, showing ground truth (GT) values (top row) along with the least-squares (LSQ) outputs (middle row) and hierarchical Bayesian modelling (HBM) outputs (bottom row). LSQ output maps tend to show more noise than the HBM output maps. **(A)**. DKI model, showing the apparent diffusion coefficient (D) and kurtosis (K) maps. **(B)**. FEXI AXR model, showing the apparent diffusion coefficient (D), filter efficiency (*σ*) and apparent exchange rate (AXR) maps.

**FIGURE 3.**
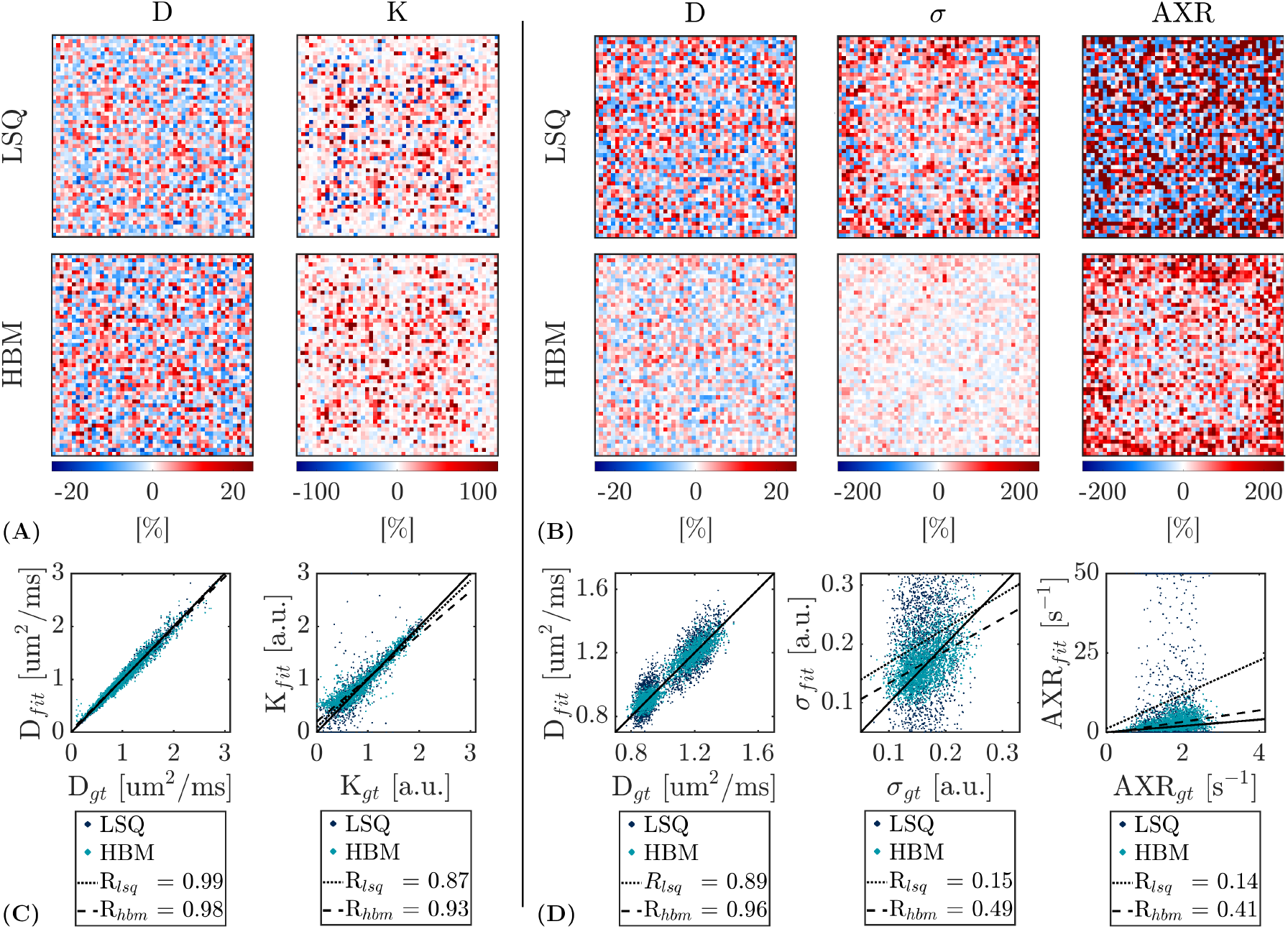
Error metrics for the least-squares (LSQ) and hierarchical Bayesian modelling (HBM) outputs in simulated data. The HBM outputs tend to show reduced errors and improved correlations with ground truth values relative to the LSQ outputs. **(A)**. Percent error maps are shown for the apparent diffusion coefficient (D) and kurtosis (K) parameters of the DKI model. **(B)**. Percent error maps for the apparent diffusion coefficient (D), filter efficiency (*σ*) and apparent exchange rate (AXR) parameters of the FEXI AXR model. **(C)**. Correlations with ground truth (GT) values for the LSQ- and HBM-derived DKI model parameters. **(D)**. Correlations with ground truth values for the LSQ- and HBM-derived FEXI AXR model parameters.

The RMSE (Table 2) was also improved using the HBM approach: it decreased by 25% for kurtosis (from RMSE_*lsq*_ = 0.20 to RMSE_*hbm*_ = 0.15) and by 89% for AXR (from RMSE_*lsq*_ = 19.2 s^−1^ to RMSE_*hbm*_ = 2.1 s^−1^). Bias was minimal in the DKI model for the LSQ and HBM outputs of both parameters; for the FEXI AXR model, the HBM fit reduced bias in the AXR estimation by 88% (from bias_*lsq*_ = 8.5 s^−1^ to bias_*hbm*_ = 1.0 s^−1^).

Minimal or no improvements were seen in the HBM diffusivity estimates of either model as it is typically a stable parameter to fit, even when using LSQ methods.

### 3.2 Contrast-to-noise

With the HBM method, CNR improved in the simulated data by a factor of 1.2 for kurtosis (from CNR_*lsq*_ = 0.6 to CNR_*hbm*_ = 0.7) and by a factor of 6 for AXR (from CNR_*lsq*_ = 0.1 to CNR_*hbm*_ = 0.6) (Table 2). This improvement was most visually striking in the AXR parameter maps (Figure 2B), where ROI boundaries were obscured by high noise in the LSQ output maps but were well resolved in the HBM output maps.

### 3.3 Noise sensitivity of regional priors

Figure 4 shows the ground truth distribution of parameter values, along with the parameter distributions obtained from the LSQ and HBM outputs and the HBM prior distributions (averaged over the MCMC chain after the burn-in period). For the DKI model (Figure 4A), the LSQ and HBM outputs and the HBM priors were well-matched to the ground truth across SNR levels. For the FEXI AXR model (Figure 4B), the HBM priors became less informative at lower SNRs, displaying wide, flat profiles; however, even this weak influence helped resolve the WM/GM ROIs, although biases were introduced. LSQ AXR outputs provided no contrast between the separate ROIs at low SNRs, with parameter estimates clustered at the bounds for both ROIs. The information content of the HBM priors improved at higher SNRs, more closely resembling the ground truth distributions.

**FIGURE 4.**
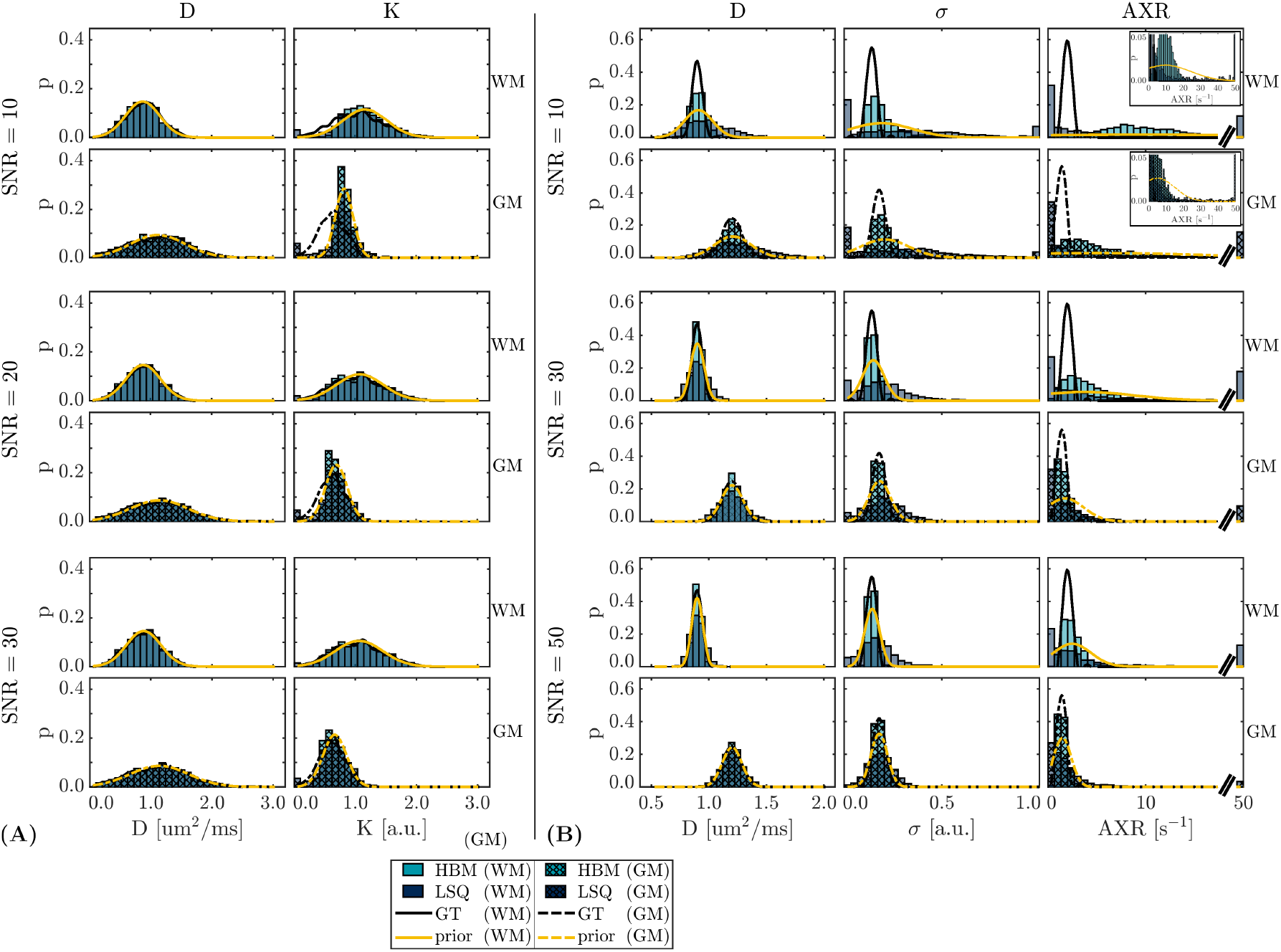
Impact of SNR on parameters fitted using least-squares (LSQ) (dark blue) and hierarchical Bayesian modelling (HBM) (teal) approaches. Block-filled bars show histograms of voxelwise parameter estimates from the simulated ‘white matter’ (WM) region; hatch-filled bars show distributions from the simulated ‘grey matter’ (GM) region. Ground truth distributions are indicated in black; HBM priors are shown in yellow. **(A)**. DKI model, showing the apparent diffusion coefficient (D) and kurtosis (K) parameter distributions. **(B)**. FEXI AXR model, showing the apparent diffusion coefficient (D), filter efficiency (*σ*) and apparent exchange rate (AXR) parameter distributions. Inset figures for SNR = 10 are re-scaled to better show the prior distributions.

### 3.4 Healthy volunteer data

Figure 5 shows parameter maps for the healthy volunteer DKI data, estimated using the LSQ method and the HBM method with *k* = 2 regional priors. A reduction in the number of extreme fits in the kurtosis estimation was shown using the HBM approach (Figure 5, inset), in accordance with the simulated data (Figure 2A). A reduction in diffusivity was observed near the ventricles in the HBM-derived maps.

**FIGURE 5.**
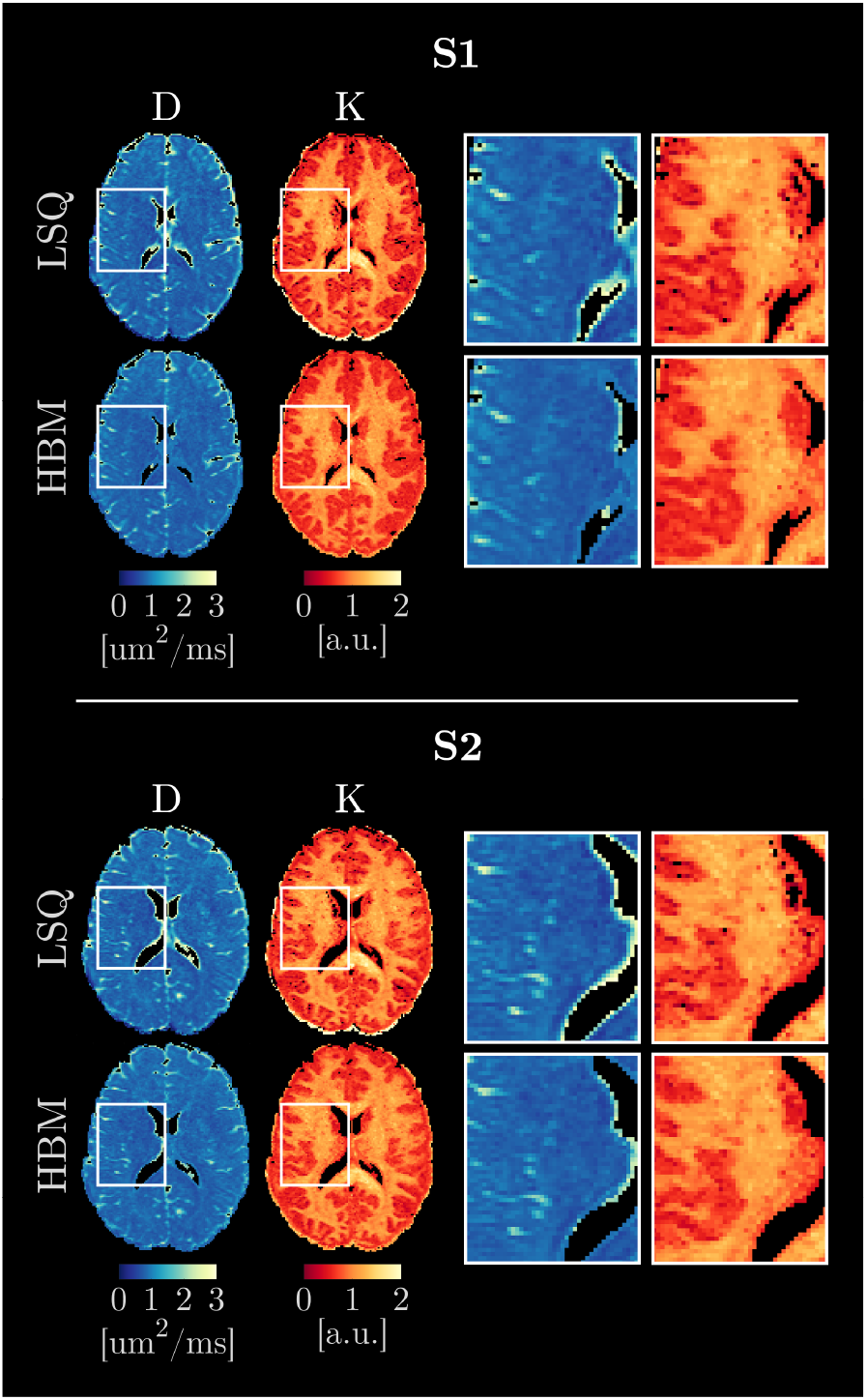
Diffusion kurtosis imaging (DKI) parameter maps for the two test cases (top panel, subject S1; bottom panel, subject S2), showing the least-squares (LSQ) and hierarchical Bayesian modelling (HBM) (with *k* = 2 regional priors: white matter and grey matter) outputs. Inset images highlight regions of reduced diffusivity (D) around the ventricles in the HBM maps (indicating a potential a reduction in CSF contamination) and reduced noise in the HBM kurtosis (K) maps.

### 3.5 Exemplar cSVD subject data

AXR parameter maps of the two exemplar cSVD subjects are shown in Figure 6 for the LSQ approach and the HBM approach with *k* = 3 regional priors; other parameters are shown for each subject respectively in Figures S1.1 and S1.2. Noise was substantially reduced in the HBM parameter maps, revealing an increase in AXR in WMH relative to both WM and GM; by comparison, high noise levels in the LSQ maps obscured any differences in the WMH AXR.

**FIGURE 6.**
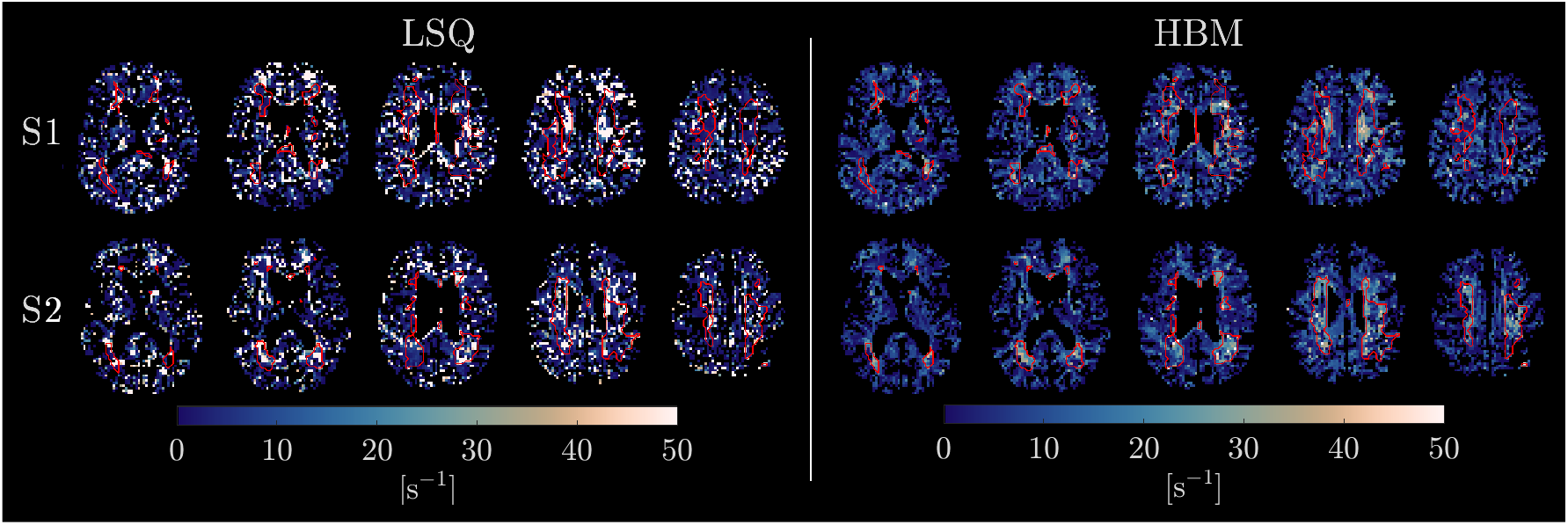
Apparent exchange rate (AXR) parameter maps for cerebral small vessel disease (cSVD) subject S1 (top row) and subject S2 (bottom row), derived using least-squares (LSQ) and hierarchical Bayesian modelling (HBM) (with *k* = 3 regional priors: white matter, grey matter, and white matter hyperintensities) approaches. Noise is substantially reduced in the HBM parameter maps, revealing regions of increased AXR within cSVD white matter hyperintensities (outlined in red) that are not visible in the LSQ maps.

Box plots of parameter estimates in WM, GM and WMH are shown in Figure 7A-B for each of the different prior configurations (*k* = 1, 2, 3). Without a separate prior for WMH (*k* = 2), median WM and WMH values were similar, with 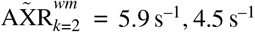 and 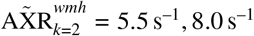 for subjects S1, S2 respectively. When using a separate prior for WMH (*k* = 3), a substantial increase in the median WMH AXR compared to the median WM AXR was observed: 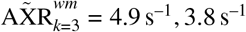 and 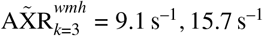 for subjects S1, S2 respectively. Median WMH AXR values estimated using the LSQ approach were considerably lower at 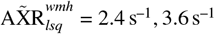 for subjects S1, S2 (with 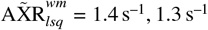), but showed a wider distribution across voxels than all of the HBM-derived estimates.

**FIGURE 7.**
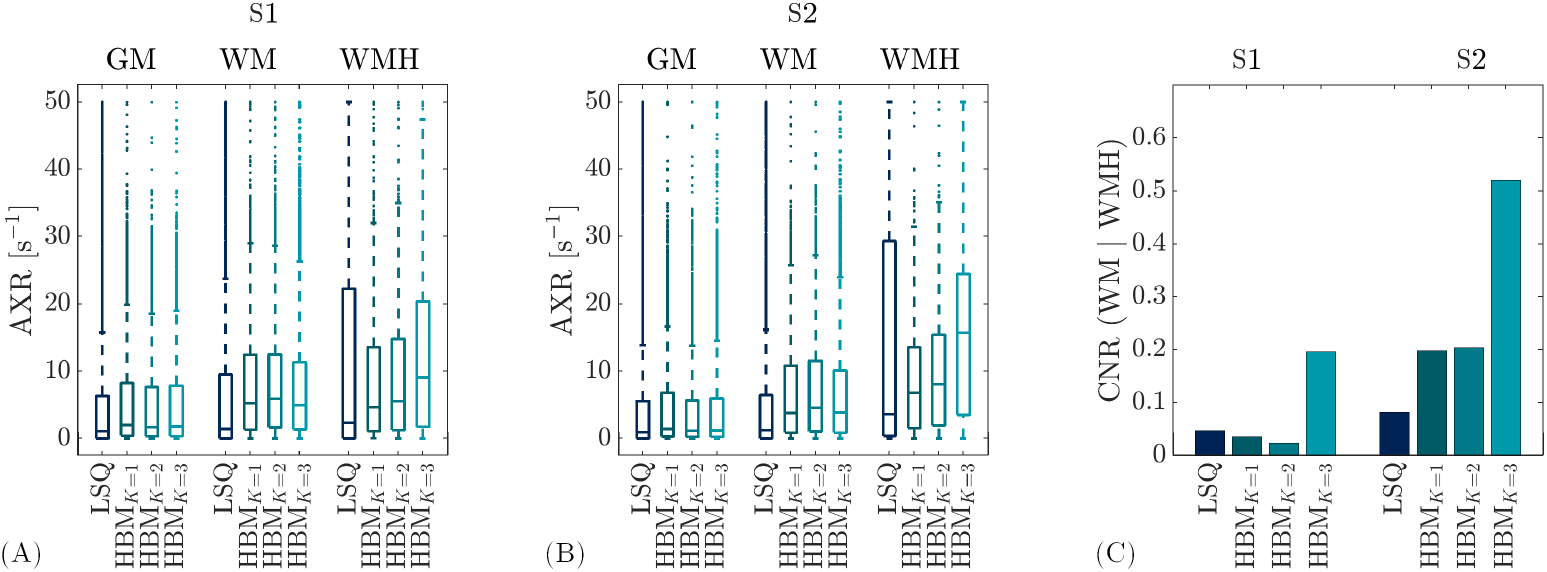
Impact of different regional priors in cerebral small vessel disease (cSVD) subjects. **(A)**. Boxplots of voxelwise apparent exchange rate (AXR) estimates are shown for cSVD subject S1 in grey matter (GM), white matter (WM) and white matter hyperintensities (WMH) for the least squares (LSQ) and hierarchical Bayesian modelling (HBM) approaches; the number of regional priors used for the HBM approach is indicated by the subscript, where *k* = 1 indicates one global prior, *k* = 2 indicates two priors (WM/GM), and *k* = 3 indicates three priors (WM/GM/WMH). Using a separate prior for WMH (i.e. *k* = 3) leads to a substantial increase in the median AXR in WMH, while median WMH AXR values for *k* = 1, 2 are closer to WM AXR values. **(B)**. As for (A), but for cSVD subject S2. **(C)**. Contrast-to-noise ratio (CNR) between WM and WMH subjects for S1 and S2. The CNR is higher for *k* = 3 than for *k* = 1, 2, indicating superior discrimination of WMH AXR alterations with a separate WMH prior.

Figure 7C shows the CNR between WM and WMH for each prior configuration. The CNR was higher for *k* = 3 (CNR_*k*=3_ = 0.2, 0.5 for subjects S1, S2 respectively) than for *k* = 1, 2 (e.g. CNR_*k*=2_ = 0.02, 0.2 for subjects S1, S2), demonstrating the potential for differentiating between WM and WMH AXR values when using a separate WMH prior. The CNR between WM and WMH was minimal when using the LSQ approach (CNR_*lsq*_ = 0.05, 0.08 for subjects S1, S2).

The average SNR in the brain, estimated using the mean and SD of the six repetitions with *b*_*f*_ = 0 ms/µm^2^, *t*_*m*_ = 16 ms, *b* = 0 ms/µm^2^, was SNR = 26, 32 in subjects S1, S2 respectively.

## 4 DISCUSSION

Our generalised algorithm, which enables HBM fitting for any dMRI microstructural model and incorporates arbitrary parameter constraints and regional priors, is demonstrated here in two previously unexplored neuroimaging applications. The algorithm is implemented in both Matlab (R2024b) and Python (v3.9.20), and is publicly available at https://github.com/e-powell/matlab-bayesian and https://github.com/e-powell/dmipy-bayesian respectively; in Python, the Dmipy ^43^ software package is utilised and extended. Note that all results reported in this work used the Matlab implementation. The HBM algorithm was shown to outperform conventional LSQ approaches in both simulated and human data for the diffusion-based DKI and BBB-FEXI AXR models.

The greatest improvements were observed for the least stable parameters in the LSQ fit (kurtosis in the DKI model; AXR in the FEXI AXR model), particularly for the more complex FEXI AXR model. Complex models often suffer from parameter degeneracy - where multiple combinations of model parameters give rise to similar MRI signals - and high sensitivity to noise; consequently, voxelwise fitting techniques often return values associated with erroneous local minima in the fitting objective function. Exploiting information across a region - as is the role of the prior distributions in the HBM method - can reduce the sensitivity of a model to these effects, as the prior information increases the likelihood of selecting parameter combinations more similar to the rest of the region.

This is exemplified in the DKI model (Figure 2A and Figure 5), where small areas of noise in the LSQ-derived kurtosis maps - which are demonstrably fitting artefacts in the simulated data - were removed in the HBM maps. Areas of high diffusivity around the ventricles in the in vivo LSQ maps were also reduced in the HBM maps, potentially indicating a reduction in partial volume effects arising from CSF contamination by the HBM method.

The benefits of the HBM approach were most apparent for the FEXI AXR model (Figure 2B, Figure 6), likely reflecting the higher noise susceptibility (and subsequently noisier LSQ fits) of FEXI AXR model parameters relative to DKI model parameters. Experiments with simulated data in this work provide confidence that, for the noise levels expected in human data ^6^, HBM can still infer the true FEXI AXR model parameter values, as demonstrated by the error maps in Figure 3. The cSVD case studies best exemplify the HBM advantages: a clear contrast between WMH AXR and surrounding brain tissue AXR values was revealed in HBM maps, while noise in the LSQ maps obscured any regional variation (Figure 6). The AXR was generally higher in WMH (although this was variable) than in surrounding tissues, suggesting increased BBB damage; this has been reported in studies using dynamic contrast-enhanced MRI ^44^. As this work was designed to introduce and demonstrate proof-of-principle of the HBM algorithm, a detailed interpretation of the observed WMH AXR values in relation to underlying pathophysiology is out of scope here; however, it is noted that cSVD is a diffuse and heterogeneous disease process with multiple factors contributing to small vessel and BBB damage ^45,46^, potentially explaining the AXR variability within WMH. Previous studies have demonstrated altered BBB water exchange in cSVD subjects using a diffusion-prepared pseudo-continuous arterial spin labelling sequence ^47,48^, although a decrease in water exchange rate was observed while an increase is detected here. It may be important to note that these previous studies included genetic causes of cSVD (representing a minority of cSVD cases) in contrast to the two cSVD subjects included in this work who are older adults with vascular risk factors; careful consideration of the differences between MRI acquisition techniques and factors such as relaxation time differences (known to influence the FEXI AXR model ^4,6^) in WMH, along with increased sample sizes, are also required before drawing conclusions. However, without loss of generalisability, these data provide evidence that localised pathology can be resolved and detected using the HBM method, even in high noise conditions.

A limitation of the HBM method is that in high noise conditions it is possible that the prior (and subsequently parameter estimates derived using HBM) may tend towards the mean of the parameter range if individual voxels do not contain sufficient information for inference. Noise sensitivity analysis on the regional priors demonstrated that, for the models chosen here, some priors became heavily biased at low SNR (for example AXR at SNR = 10; Figure 4B), but maintained enough information content to resolve the separate ROIs and did not reduce parameter distributions to the mean of the parameter bounds.

A related issue under high noise conditions - and one of the primary concerns of any hierarchical method - is that regions with underlying parameter values distinct from the regional prior (e.g. lesions) may become obscured. Experiments on simulated DKI and BBB-FEXI data using just one global prior (Supporting Information S2) confirmed that the two distinct ‘WM’ and ‘GM’ ROIs were still resolved, suggesting that, at the noise levels used here and the number of samples (voxels) within each ROI, the regional prior did not dominate the voxelwise likelihoods. However, in the cSVD patient data, using a separate regional prior for WMH substantially improved discrimination between WM and WMH AXR estimates (Figure 7). The unbalanced ROI sample sizes between WM (*∼* 10, 000 voxels) and WMH (*∼* 1, 200 voxels) in combination with the large difference between underlying WM/WMH AXR values (as observed for *k* = 3 regional priors) could explain this finding: a prior combining two distinct ROIs will be dominated by the larger ROI (i.e. WM here) as voxels in the smaller ROI (i.e. WMH here) will be treated as outliers by the prior, such that greater biases will be incurred in the smaller ROI for larger differences between underlying regional parameter means. For example, AXR prior means (Equation 9) were substantially different for WM and WMH when using *k* = 3 regional priors (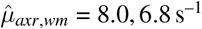 and 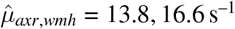 for subjects S1, S2 respectively), while the WM AXR prior mean for *k* = 2 (i.e. no WMH regional prior) was skewed towards the WM prior mean observed for 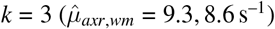. This supports the suggestion that, when WM and WMH are combined under one regional prior, the likelihood of selecting parameter combinations that reflect the true WMH distribution is reduced.

The above considerations suggest a requirement for identification of localised pathology before applying the HBM method, particularly in the context of noisier data with more complex models. This could be addressed by developing an adaptive hierarchical model that selects the number of ROIs during inference to determine the appropriate number of regional priors. Identifying additional independent ROIs - i.e. spatially-localised voxel clusters with similar parameter values - and updating voxel ROI membership accordingly at each MCMC step may improve parameter estimation in small distinct regions. However, small ROIs are more susceptible to noise, and hence the trade-off between regional prior size and noise bias needs to be carefully managed. Simulations could help elucidate any such sample size dependencies. An adaptive model could additionally remove the need for obtaining tissue segmentations from separate MRI acquisitions, as was required here for the cSVD data.

Another avenue of future work is to develop non-Gaussian priors. It was assumed here that a Gaussian prior was suitable for all model parameters; however, particularly for models that include fibre orientation parameters (e.g. NODDI ^1^), a uniform prior may instead be more appropriate. A further assumption of the HBM method is that signal noise is Gaussian, but this may not be suitable for very low signal voxels (SNR ≲ 2) ^49^, where the noise distribution becomes Rician. Parallel imaging further alters the noise characteristics ^50^, which may show non-central chi or Rayleigh distribution properties; more advanced noise reduction methods, for example denoising complex channel data prior to image reconstruction ^51^, could be included to reduce these effects. Finally, it would be prudent in future work to perform comparisons between different MCMC model fitting methods (e.g. Harms et al. ^52^).

## 5 CONCLUSIONS

Our generalised HBM and MCMC algorithm, which enables parameter inference for dMRI signal models with arbitrary parameter constraints, is shown to improve the accuracy, precision, CNR, and detection of localised pathology over the conventional voxelwise LSQ approach, providing a new option for reducing erroneous fits in complex dMRI models. The algorithm is open-source, and implemented in Matlab (https://github.com/e-powell/matlab-bayesian) and Python (https://github.com/e-powell/dmipy-bayesian).

## ACKNOWLEDGMENTS

Thanks to the radiographers at Manchester University imaging facilities for facilitating the cSVD data acquisition. Thanks to Dr David Higgins of Philips Healthcare MR Clinical Science for supporting this work.

## CONFLICTS OF INTEREST

GJMP receives salary from, is a director of, and a shareholder in Bioxydyn Limited, a company with an interest in imaging biomarkers. GJMP is also a director of and a shareholder in Queen Square Analytics Limited and is a director of and a shareholder in Quantitative Imaging Limited, companies with an interest in imaging biomarkers. GJMP has current grant funding from Eli Lilly and GSK, and received other support from Philips and Siemens Healthineers.

## DATA AVAILABILITY

The data that support the findings of this study are available on request from the corresponding author.The data are not publicly available due to privacy or ethical restrictions.

**APPENDIX**

## A FULL DERIVATION OF THE GENERAL HIERARCHICAL BAYESIAN MICROSTRUCTURE MODEL

In this section the HBM, and MCMC algorithm used for inference, are motivated and introduced in full.

### A.1 General microstructure model

Consider a general multi-compartment model of *C* compartments, with an underlying microstructure-related parameter set ***θ***. For notational convenience, model parameters ***θ*** are grouped by type as:

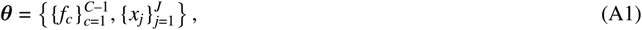

Where *f*_*c*_ denotes compartment signal fractions and ***X***_***j***_ all other parameters (e.g. diffusivities, orientations, radii). Assuming relaxation times are fixed across compartments, the signal fractions sum to one 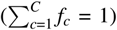, meaning that *f*_*c*_ is not a free parameter but fixed as 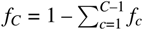.

The general multi-compartment model comprises a function,denoted *g*, that relates the underlying tissue-related parameters 0 and dMRI acquisition parameters (e.g.b-value, gradient orientations) to the measured signal intensity.For a dMRI acquisition with N diffusion-weightings, the set of acquisition parameters is defined as ***t*** = *[t1*, …, *t*_*N*_]^*T*^ and the measured signal in voxel *i* as ***S***_***i***_; = [S1, …,SN]^*T*^.The signal equation across a whole dMRI acquisition can thus be written:

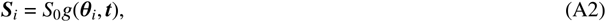

Where *S*_0_ is the signal intensity without diffusion weighting and ***θ***; are the model parameters in voxel *i*. The RHS of Equation A2 is a column vector where the nth element, S_***0***_*g*(***θ***_***i***_;,*t*_*n*_), is the model-predicted signal for diffusion weighting *t*_*n*_ given model parameters ***θ***_***i***_;.

The experimentally-measured signal for voxel *i* in the presence of Gaussian noise is hence modelled as:

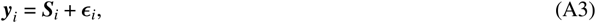

where **y** = [y_***1***_, y_2_, …, ***Y***_***N***_]^*T*^ is the measured signal and 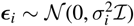 is the noise, and where *N(****µ*, Σ**) is a multivariate normal distribution with mean ***µ*** and covariance matrix **Σ**, with *σ*_*i*_ a scalar giving the noise standard deviation in voxel i and **ℐ** the N x N identity matrix.

The likelihood for the measurement in voxel *i* is:

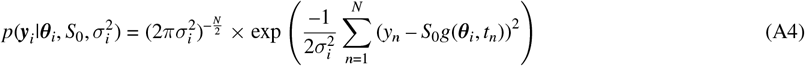

Orton et al. ^16^ demonstrated that the ‘nuisance parameters’ *S*_0_ and 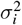 can be marginalised out from Equation A4 to give the following marginalised likelihood:

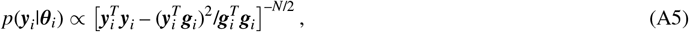

where ***g***_*i*_ = [*g*(***θ***_*i*_, *t*_1_), …, *g*(***θ***_*i*_, *t*_*N*_)] are the model-predicted signals for voxel *i*.

### A.2 Bayesian shrinkage priors

Orton et al. ^16^ used a multivariate Gaussian Bayesian shrinkage prior (BSP) on the IVIM model parameters over a single user-defined ROI, defined as:

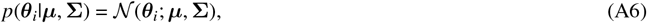

where ***µ*** and **Σ** are estimated from the data, and effectively encode the mean and covariance respectively of all model parameters across the ROI.

The method of Orton et al. ^16^ is extended here to account for multiple ROIs. Given *K* ROIs, denoted *k* = 1, …, *K*, each voxel is a member of exactly one ROI. Each ROI *k* has its own Gaussian BSP distribution. For voxel *i* in ROI *k*, the prior on the parameters ***θ***_*i*_ is then:

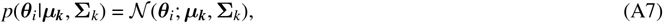

where ***µ***_***k***_ is a vector whose elements encode the prior means of the parameters in ROI *k*, **Σ**_*k*_ is their covariance and *N* (***θ***; ***µ*, Σ**) denotes the multivariate normal probability density function (PDF). Again, it is emphasised that ***µ***_*k*_ and **Σ**_*k*_ are estimated from the data.

To generalise from Orton et al.’s ^16^ two-compartment model to an arbitrary multi-compartment model, all signal fractions must to sum to one. This is enforced here (following Harms et al. ^52^) by modifying the priors to:

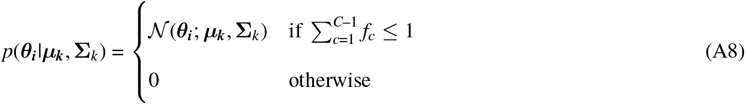

To complete the model, hyper-priors are defined on all ***µ***_*k*_’s and **Σ**_*k*_’s as non-informative Jeffrey’s priors:

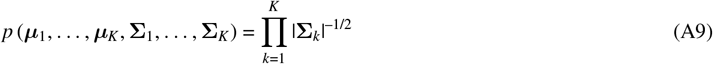

### A.3 Posterior distributions

Each ROI has an independent posterior distribution. If ROI *k* comprises *I*_*k*_ voxels, its posterior distribution can be written as ^16^:

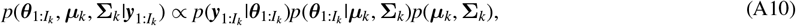

where 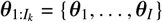 are the parameters and 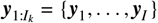 the dMRI data for all *I*_*k*_ voxels in the ROI. Substituting in Equations A5, A7 and A9 gives:

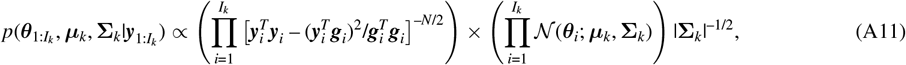

from which samples can be drawn with an MCMC algorithm (see Appendix A.5).

### A.4 Parameter transforms

Microstructure model fitting needs to enforce physically reasonable minimum and maximum values of parameters; for example, diffusivities need to be positive. Here, the transforms used by Orton et al. ^16^ are generalised to enable arbitrary minimum and maximum constraints. For a single parameter *θ*, its transform is defined as:

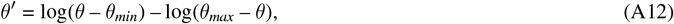

which maps the interval (*θ*_*min*_, *θ*_*max*_) to ℝ. By defining the Bayesian prior on the transformed parameter *θ*^′^, *θ* is therefore constrained between *θ*_*min*_ and *θ*_*max*_.

### A.5 MCMC algorithm

The MCMC algorithm is derived here, and given as pseudocode in Algorithm 1.

#### A.5.1 ROI-wide parameters

Following Orton et al. ^16^, the MCMC updates for the ROI-wide prior parameters ***µ***_1:*K*_ and **Σ**_1:*K*_ are Gibbs moves. The conditional distributions are (up to proportionality):

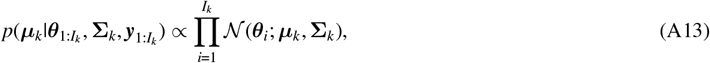

where 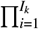 denotes the product over all voxels in ROI *k*. By rearranging the multivariate normal PDF so that ***µ***_*k*_ is the variable, it is given that:

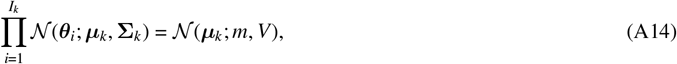

where

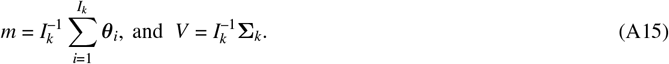

The MCMC updates are therefore sampled as:

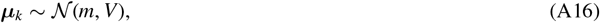

where *N* (*m, V*) is a multivariate normal distribution with mean *m* and covariance *V*. Following the same steps for **Σ**_*k*_ (full details in Orton et al. ^16^) gives the MCMC update:

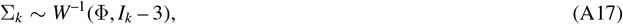

where

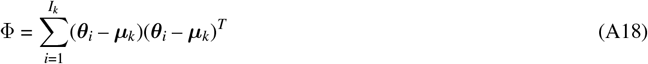

and *W*^−1^ is the inverse-Wishart distribution.

#### A.5.2 Voxelwise parameters

For the non signal fraction voxelwise parameters the posterior distribution for voxel *i* in ROI *k*, up to proportionality, is:

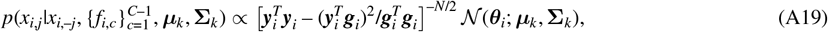

where *x*_*i*,*j*_ is the *j*th parameter value, *x*_*i*,–*j*_ = {*x*_*i*,1_, …, *x*_*i*,*j*–1_, *x*_*i*,*j*+1_, …, *x*_*i*,*J*_} denotes the set of all non signal fraction parameters except *x*_*i*,*j*_, and 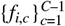 are the signal fractions.

As in Orton et al. ^16^, this is sampled with a Metropolis-Hastings algorithm. Proposed parameters, 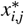, are first sampled from Gaussian distributions as:

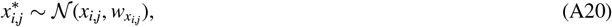

where *x*_*i*,*j*_ is the current parameter value and 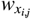 the proposal distribution variance, which should reflect the parameter scale and can be tuned for optimal algorithm performance.

The acceptance probability utilises the ratio of the posterior distributions for *x*_*i*,*j*_ and 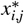:

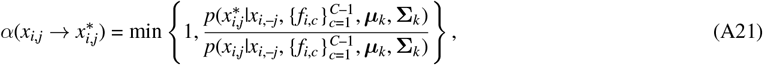

where the numerator and denominator are calculated using Equation A19.

MCMC moves for the signal fractions are the same, except that the posterior distributions now contain the terms enforcing 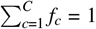:

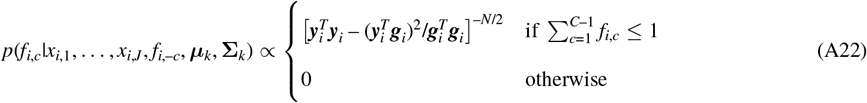

where *f*_*i*,–*c*_ = {*f*_*i*,1_, …, *f*_*i*,*c*–1_, *f*_*i*,*c*+1_, …, *f*_*i*,*C*_} are the other signal fractions apart from *f*_*i*,*c*_. Again, proposed values are sampled as:

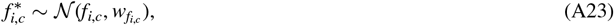

where *f*_*i*,*c*_ is the current signal fraction. The acceptance probabilities are:

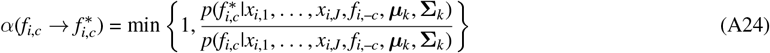

#### A.5.3 Metropolis-Hastings acceptance ratio

Metropolis-Hastings jumping variances are tuned during the first half of the burn-in period to achieve an acceptance ratio that efficiently samples the posterior distribution. Variances are adjusted voxelwise for all compartment signal fractions, 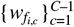, and other parameters, 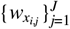. The tuning strategy of Orton et al. ^16^ is extended here by applying the following update rule at every *n*_*s*_ MCMC steps:

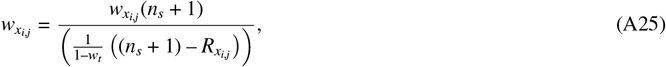

where 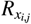 is the number of accepted updates for parameter *x*_*i*,*j*_ during the previous *n*_*s*_ steps, and *w*_*t*_ is the target acceptance rate. Compartment signal fractions are updated similarly:

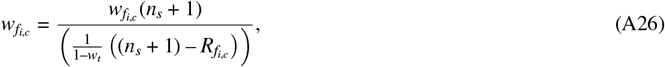

The target acceptance rate is set to *w*_*t*_ = 0.25^53^, adjusting the jumping variances to aim for an acceptance rate of 25%.

## Algorithm 1

Pseudocode for hierarchical Bayesian model fitting of a general microstructural model and the MCMC algorithm used for inference.

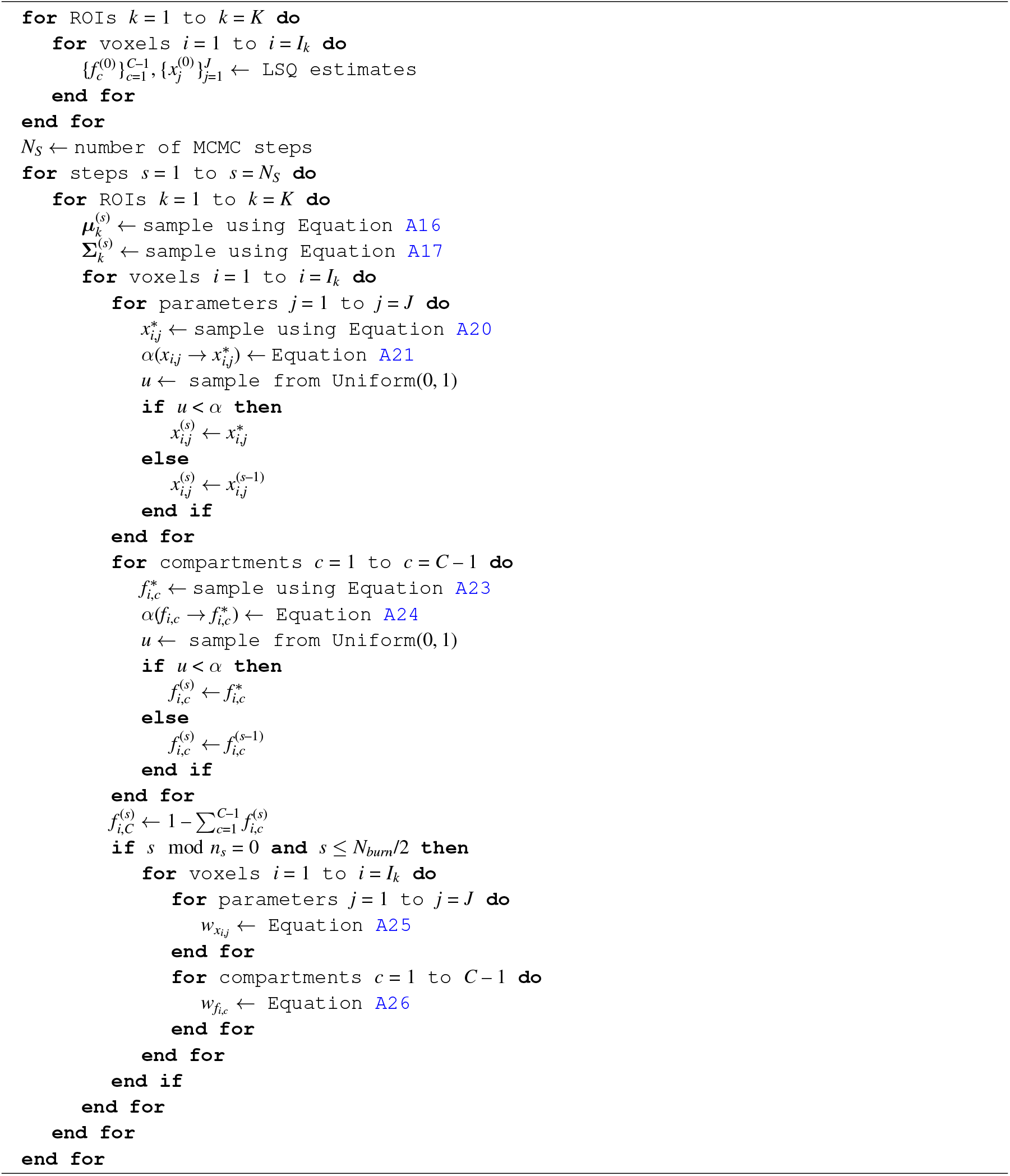

## SUPPORTING FIGURES AND TABLES

### S1 CEREBRAL SMALL VESSEL DISEASE PATIENTS

All parameters of the BBB-FEXI AXR model are shown here for the two exemplar cSVD subjects (Figures S1.1 and S1.2 respectively), both for the LSQ approach and for the HBM approach with *k* = 3 regional priors. Noise was substantially reduced in the HBM parameter maps, revealing an increase in AXR in WMH relative to WM; by comparison, high noise levels in the LSQ maps obscured any alterations in the WMH AXR.

**FIGURE S1.1.**
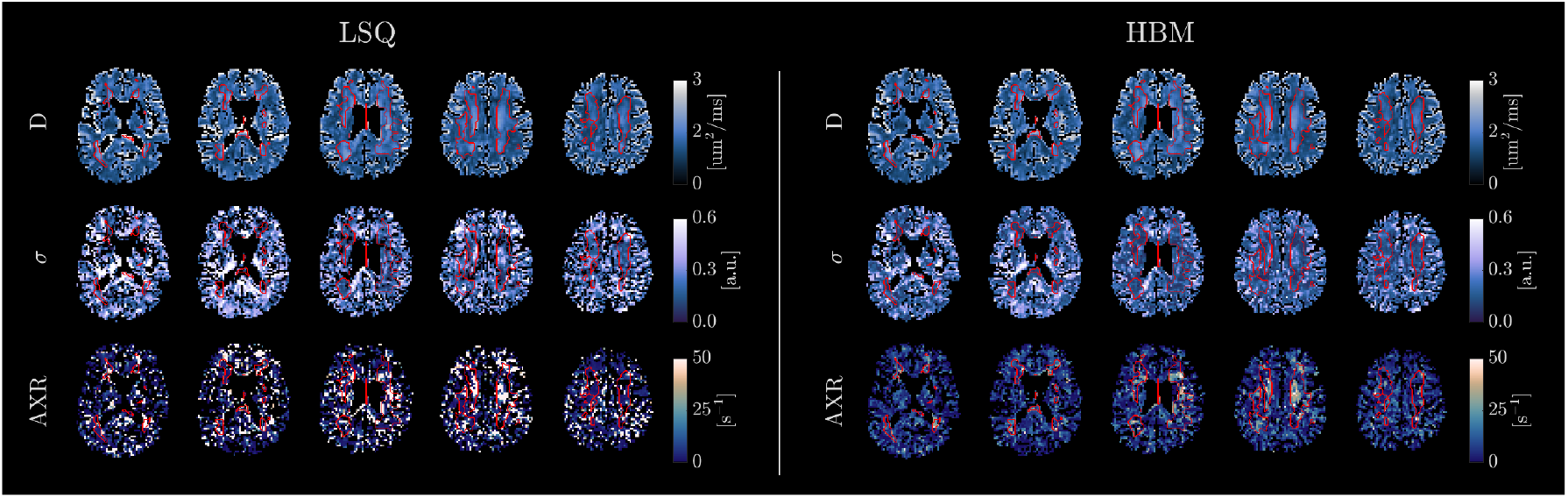
Cerebral small vessel disease subject S1. Apparent diffusion coefficient (D), filter exchange (*σ*) and apparent exchange rate (AXR) parameter maps, derived using least-squares (LSQ) and hierarchical Bayesian modelling (HBM) (with *k* = 3 regional priors: white matter, grey matter, and white matter hyperintensity) approaches. Noise is substantially reduced in the HBM parameter maps, revealing regions of increased AXR within white matter hyperintensities (outlined in red) that are not visible in the LSQ maps.

**FIGURE S1.2.**
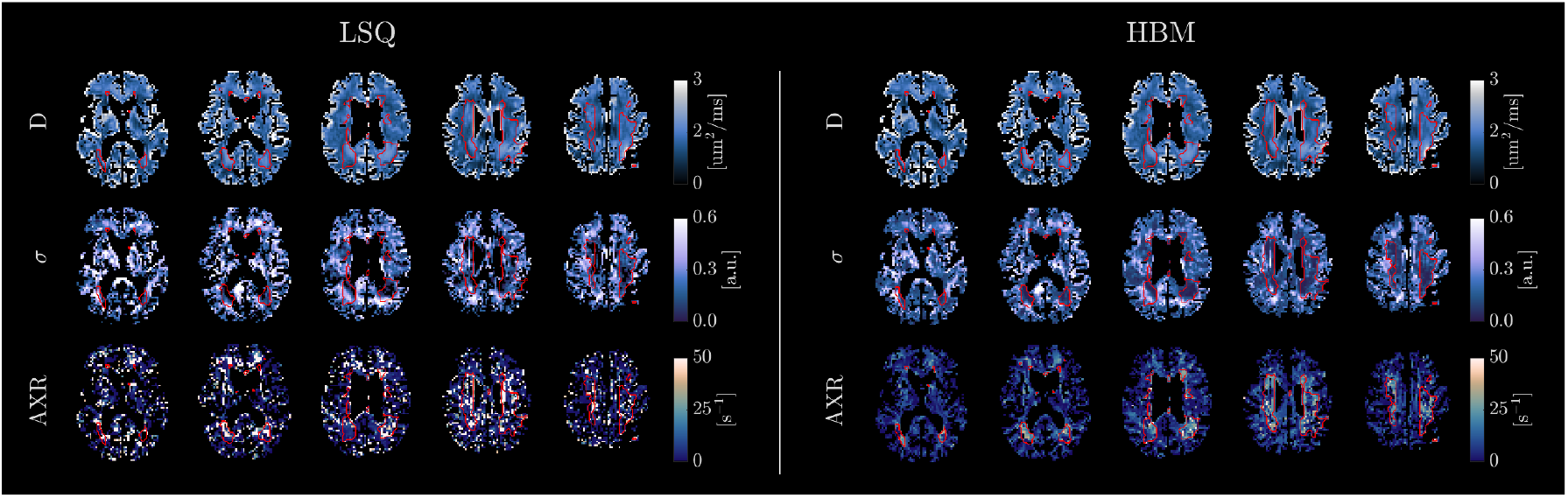
Cerebral small vessel disease subject S2. Apparent diffusion coefficient (D), filter exchange (*σ*) and apparent exchange rate (AXR) parameter maps, derived using least-squares (LSQ) and hierarchical Bayesian modelling (HBM) (with *k* = 3 regional priors: white matter, grey matter, and white matter hyperintensity) approaches. Noise is substantially reduced in the HBM parameter maps, revealing regions of increased AXR within white matter hyperintensities (outlined in red) that are not visible in the LSQ maps.

### S2 HBM WITH ONE GLOBAL PRIOR (SIMULATIONS)

The DKI and AXR models were fitted to the data generated in Sections 2.3.1 and 2.3.2 respectively, using the HBM approach with *k* = 1 global prior (cf. the original implementation using *k* = 2 regional priors, one for each simulated ‘WM’ and ‘GM’ ROI). Parameter maps for these HBM outputs are shown in Figure S2.1. Error maps and statistical analyses are provided in Figure S2.2 and Table S2 respectively.

For the DKI model, both ROIs remained well defined (Figure S2.1) with good parameter accuracy (Figure S2.2, Table S2) using a single global prior. For the FEXI AXR model, there was visibly less contrast between ROIs in the AXR parameter map using one global prior (CNR_*hbm*(1)_ = 0.5 s^−1^ and CNR_*hbm*(2)_ = 0.6 s^−1^, for *k* = 1, 2 priors respectively); however, parameter estimates were less biased (bias_*hbm*(1)_ = 0.6 s^−1^ and bias_*hbm*(2)_ = 1.0 s^−1^, for *k* = 1, 2 priors respectively).

**FIGURE S1.2.**
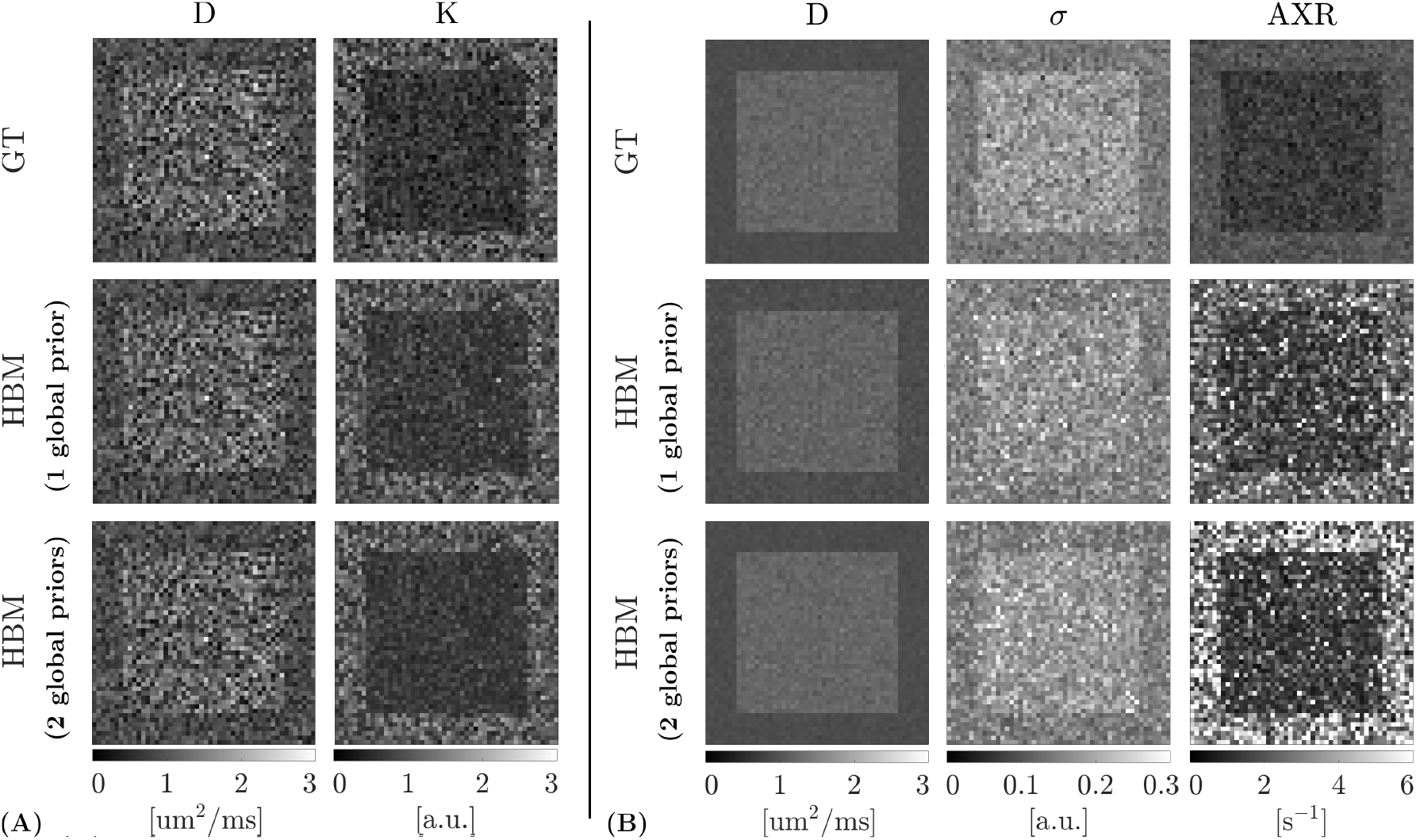
Parameter maps for the simulated data, showing ground truth (GT) values (top row) along with the hierarchical Bayesian modelling (HBM) outputs using *k* = 1 global prior (middle row). The original HBM estimates using *k* = 2 regional priors are replicated for comparison (bottom row; from Figure 2B). Both ROIs are still resolved for all parameters using *k* = 1 global prior. **(A)**. DKI model, showing the apparent diffusion coefficient (D) and kurtosis (K) maps. **(B)**. FEXI AXR model, showing the apparent diffusion coefficient (D), filter efficiency (*σ*) and apparent exchange rate (AXR) maps.

**FIGURE S2.2.**
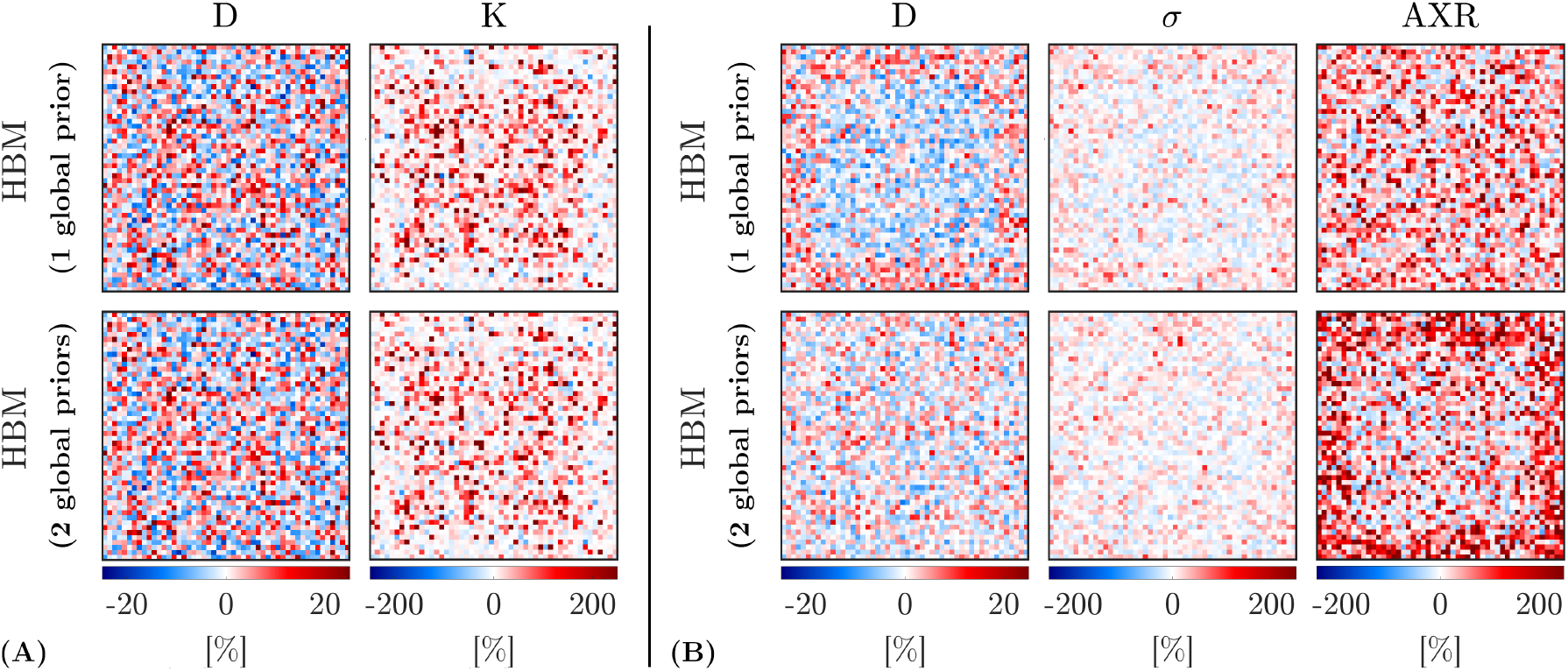
Error maps for the hierarchical Bayesian modelling (HBM) outputs in simulated data estimated using *k* = 1 global prior (top row); errors maps using the original HBM approach with *k* = 2 regional priors are replicated for comparison (bottom row; from Figure 3B). Error maps are comparable for *k* = 1 and *k* = 2 for most parameters, although some parameters show differences between approaches. **(A)**. Percent error maps are shown for the apparent diffusion coefficient (D) and kurtosis (K) parameters of the DKI model. **(B)**. Percent error maps for the apparent diffusion coefficient (D), filter efficiency (*σ*) and apparent exchange rate (AXR) parameters of the FEXI AXR model.

**TABLE S2.** Error metrics for the simulated DKI (SNR = 20) and FEXI AXR (SNR = 30) data, estimated using *k* = 1 global prior for the hierarchical Bayesian model (HBM) fitting approach (denoted ‘HBM (1)’). Values from the original *k* = 2 prior implementation are replicated here for comparison (denoted ‘HBM (2)’; from Table 2).

|  |  | DKI model |  | AXR model |  |  |
| --- | --- | --- | --- | --- | --- | --- |
| | | $D$<br>[$\mu\text{m}^2/\text{ms}$] | $K$<br>[a.u.] | $D$<br>[$\mu\text{m}^2/\text{ms}$] | $\sigma$<br>[a.u.] | AXR<br>[$\text{s}^{-1}$] |
| RMSE | HBM (1) | $8.1 \times 10^{-2}$ | 0.17 | $5.7 \times 10^{-2}$ | $3.8 \times 10^{-2}$ | 1.3 |
| | HBM (2) | $7.9 \times 10^{-2}$ | 0.15 | $4.6 \times 10^{-2}$ | $3.5 \times 10^{-2}$ | 2.1 |
| Bias | HBM (1) | $-0.1 \times 10^{-2}$ | 0.06 | $-0.4 \times 10^{-2}$ | $0.4 \times 10^{-2}$ | 0.6 |
| | HBM (2) | $0.2 \times 10^{-2}$ | 0.05 | $0.1 \times 10^{-2}$ | $0.6 \times 10^{-2}$ | 1.0 |
| CNR | GT | 0.3 | 0.7 | 2.6 | 0.8 | 1.0 |
|  | HBM (1) | 0.3 | 0.6 | 1.8 | 0.4 | 0.5 |
|  | HBM (2) | 0.3 | 0.7 | 2.8 | 0.6 | 0.6 |

**Abbreviations:** AXR, apparent exchange rate; CNR, contrast-to-noise ratio; DKI, diffusion kurtosis imaging; RMSE, root mean squared error.

There is likely to be some interplay between the chosen number of regional priors (*k*), the number of samples (voxels) in each prior ROI, and the underlying regional parameter means that will influence the ability of the HBM method to resolve distinct regions accurately.

## Abbreviations

ADC: apparent diffusion coefficient
AXR: apparent exchange rate
BSP: Bayesian shrinkage prior
BBB: blood-brain barrier
CNR: contrast-to-noise ratio
cSVD: cerebral small vessel disease
DKI: diffusion kurtosis imaging
DTI: diffusion tensor imaging
dMRI: diffusion MRI
FA: fractional anisotropy
FEXI: filter exchange imaging
FLAIR: fluid attenuated inversion recovery
IQR: interquartile range
HBM: hierarchical Bayesian modelling
GM: grey matter
HCP: Human Conuectome Project
IVIM: intravoxel incoherent motion
LSQ: least-squares
MCMC: Markov chain Monte Carlo
NSA: number of signal averages
PDF: probability density function
SD: standard deviation
RMSE: root mean squared error
ROI: region of interest
SNR: signal-to-noise ratio
WM: white matter
WMH: white matter hyperinteusity

## Notes

Funding Information This research was supported by the EPSRC (grants EP/S031510/l and EP/M020533/l) and the Alzheimer’s Society Heather Corrie Impact Fund (grant no. 577).

### Competing Interest Statement

The authors have declared no competing interest.

